# From resonance to chaos: modulating spatiotemporal patterns through a synthetic optogenetic oscillator

**DOI:** 10.1101/2024.03.28.586779

**Authors:** Jung Hun Park, Gábor Holló, Yolanda Schaerli

## Abstract

Oscillations are a recurrent phenomenon in biological systems across scales, including circadian clocks, metabolic oscillations and embryonic genetic oscillators. Despite their fundamental significance in biology, deciphering core principles of biological oscillators is very challenging due to the multiscale complexity of genetic networks and the difficulty in perturbing organisms *in vivo*. In this study, we tackle this challenge by re-designing the well-characterised synthetic oscillator, known as “repressilator”, in *Escherichia coli* and controlling it using optogenetics, thus introducing the “optoscillator”. When we apply periodic light pulses, the optoscillator behaves as a forced oscillator. Bacterial colonies harboring synthetic oscillators manifest oscillations as spatial ring patterns. Leveraging this feature, we systematically investigate the number, intensity and sharpness of the rings under different regimes of light exposure. By integrating experimental approaches with mathematical modeling, we show that this simple oscillatory circuit can generate complex dynamics that, depending on the external periodic forcing, are transformed into distinct spatial patterns. We report the observation of synchronisation, resonance, undertone and period doubling. Furthermore, we present evidence supporting the existence of a chaotic regime. This work highlights the intricate spatiotemporal patterns accessible by synthetic oscillators and underscores the potential of our approach in understanding the underlying principles governing biological oscillations.

## Introduction

Oscillations - rhythmic and repetitive variations in time - are occurring in all organisms and are essential for achieving self-organization and biological complexity [1]. Examples of oscillations in biology include cellular metabolic activities [2–4], cell cycle control [5, 6], physiological processes such as circadian rhythms [7], spatiotemporal pattern formation during organisms’ development [8, 9] and even population dynamics and ecology [10, 11]. Despite the ubiquitous presence of oscillations in biology, our current understanding of complex dynamic properties of oscillators – such as resonance, period-doubling and chaos – originates mainly from physics and chemistry. However, several studies indicate that these phenomena also play an important role in biological oscillators. For example, studies indirectly demonstrated the resonance phenomenon in circadian rhythm of organisms when correlating longevity or fitness to different light/dark regimes [12, 13]. Other studies have consistently shown evidence for chaos in interactions between ecological and microbial communities [14–16] and arrhythmic oscillations caused by non-linear phenomena seem to be linked to heart and neurological disorders [17, 18]. The importance of understanding the emergence of these phenomena in biology is therefore evident, however the complexity of biological systems make the experimental investigation of those properties very challenging, and consequently, the research on the intricate nature of biological oscillators remains in early stages [19, 20].

Synthetic biology offers a complementary approach to study non-linear dynamic systems. It combines principles from engineering and biology to design and construct controllable synthetic circuits with minimal interference from the host regulatory network [21]. Considered a hallmark in the field, the repressilator was the first synthetic oscillator to be implemented *in vivo*, namely in the bacterium *Escherichia coli* [22]. This circuit is a three-node negative feedback loop, where each of the three repressors inhibits the expression of the next one in the loop (TetR⊣1LacI⊣1cI⊣1TetR). While the first version suffered from irregular oscillations, improved molecular implementations with the same [23] or different repressors [24–27] led to highly robust oscillations. Since the repressilator, many other synthetic oscillatory circuits with different topologies have been established in cell-free systems, bacteria and mammalian cell cultures (reviewed in [28, 29]), providing an improved understanding of oscillatory systems in biology. They also have a big potential for applications. Examples include oscillators for cancer treatment [30], as marker of bacterial growth in the gut [31], to slow down cellular aging [32] and for investigating the importance of phenotypic variation in bacterial infections [33]. Moreover, synthetic oscillations have also been used to generate spatial patterns [34]. For example, growing *E. coli* colonies containing the improved repressilator form concentric rings of the fluorescent reporters [23]. This pattern is formed because only the cells at the colony edge are oscillating and growing, while cells in the interior are arrested in different phases of the oscillations. This simple method of analysing rings in colonies opens the possibility of studying patterns that are formed through oscillatory behaviours.

Most of previously built synthetic oscillators lost synchrony after few periods [22] or were synchronised via cell-cell communication [35][36]. On the other hand, many natural oscillators, such as the circadian clock, rely on a periodic external signal to adjust their phase and stay synchronised [12, 13] in a process known as entrainment. An oscillator positively affected by a periodic externally applied stimulus is a forced or driven oscillator, and recent efforts have employed this approach to control oscillators. Mondragon-Palomino and colleagues showed entrainment and resonance of a 2-node synthetic oscillator in *E. coli* [37] . Later, Aufinger and colleagues [20] showed period-doubling in a cell-free synthetic oscillator. In both studies, the use of a microfluidic system was essential to enable the periodic supplementation and removal of the chemical inducer in the medium. Recently, Cannarsa and colleagues [38] used optogenetics to construct a repressilator that was periodically inhibited by light, which allowed for a long lasting synchronisation of individual *E. coli* cells and detuning from their natural frequency. Contrary to chemical inducers, using light with varying intensity and period to control oscillatory behaviour confers the advantage of adjusting the level of induction in space and time with precision, without the need of refreshing the medium [39, 40].

Here, we build on previous studies to construct a forced synthetic optogenetic oscillator (we call it the “optoscillator”) in *E. coli* and investigate the emergence of complex oscillatory dynamics. We take advantage of a well-characterised light sensor [39] to apply an activating external forcing to the repressilator [23]. The precise control provided by the optogenetic system allows us to test different regimes of constant or periodic forcing on growing colonies and we then analyse the ring patterns generated from oscillations in those colonies [23, 41, 42]. By combining experiments and mathematical modeling, we show that bacteria harboring our optoscillator are capable of synchronization and we report the quantitative observation of resonance, undertone and period-doubling (period-2 and period-4). Moreover, we provide strong evidence for chaos. Our work contributes to a deeper understanding of biological oscillators and paves the way for manipulating oscillations within living organisms through synthetic biology, as well as the translation of these oscillations into intricate spatial patterns..

## Results

### Construction of a light-inducible repressilator

To construct an optogenetic oscillator, we decided to combine the blue light system from the red, green, and blue (RGB) color vision circuitry developed for *E. coli* [39, 43] with the improved version of the repressilator (pLPT107) [23]. In our design, the circuit should oscillate upon light exposure, but not display oscillations when kept in the dark. Briefly, the blue light-inducible system consists of a light sensing histidine kinase (YF1) that is active in the dark and switched off by blue light (470 nm). YF1 phosphorylates its cognate response regulator FixJ, which then activates the expression of the PhiF repressor. Thus, upon light exposure the expression of PhiF is stopped. Without the repressor, the DNA-binding domain (called T3) of the T7 split-phage RNA polymerase (RNAP) is produced to dimerize with the constitutively expressed core of the T7 RNAP, enabling it to bind to the pT3 promoter and activate the expression of the downstream gene [44].

To couple the light-inducible system to the repressilator, we opted to modify the TetR node of the repressilator to make it light-inducible. We designed a hybrid promoter (PT3lacO) which is repressed by LacI and (indirectly) activated by blue light in the presence of the blue light activation system. We first characterised the functionality of this promoter in a reporter construct by controlling the expression of mCherry with light and isopropyl *β*-d-1-thiogalactopyranoside (ITPG) – an inhibitor of LacI (Figure 1a). We grew colonies on agar plates in the presence of different intensities of light, controlled by a LED illumination tool (LITOS) [45]. After 96 h of incubation, we imaged the colonies with a fluorescence microscope and quantified the red fluorescence. We found that in absence of IPTG, there was no significant change in expression of the reporter, even with increasing light intensity. Conversely, when IPTG (1 mM) was added to the medium, mCherry expression increased proportionally with light intensity. This data shows that our hybrid promoter is (indirectly) induced by light and tightly repressed by LacI (Figure 1b).

**Figure 1:**
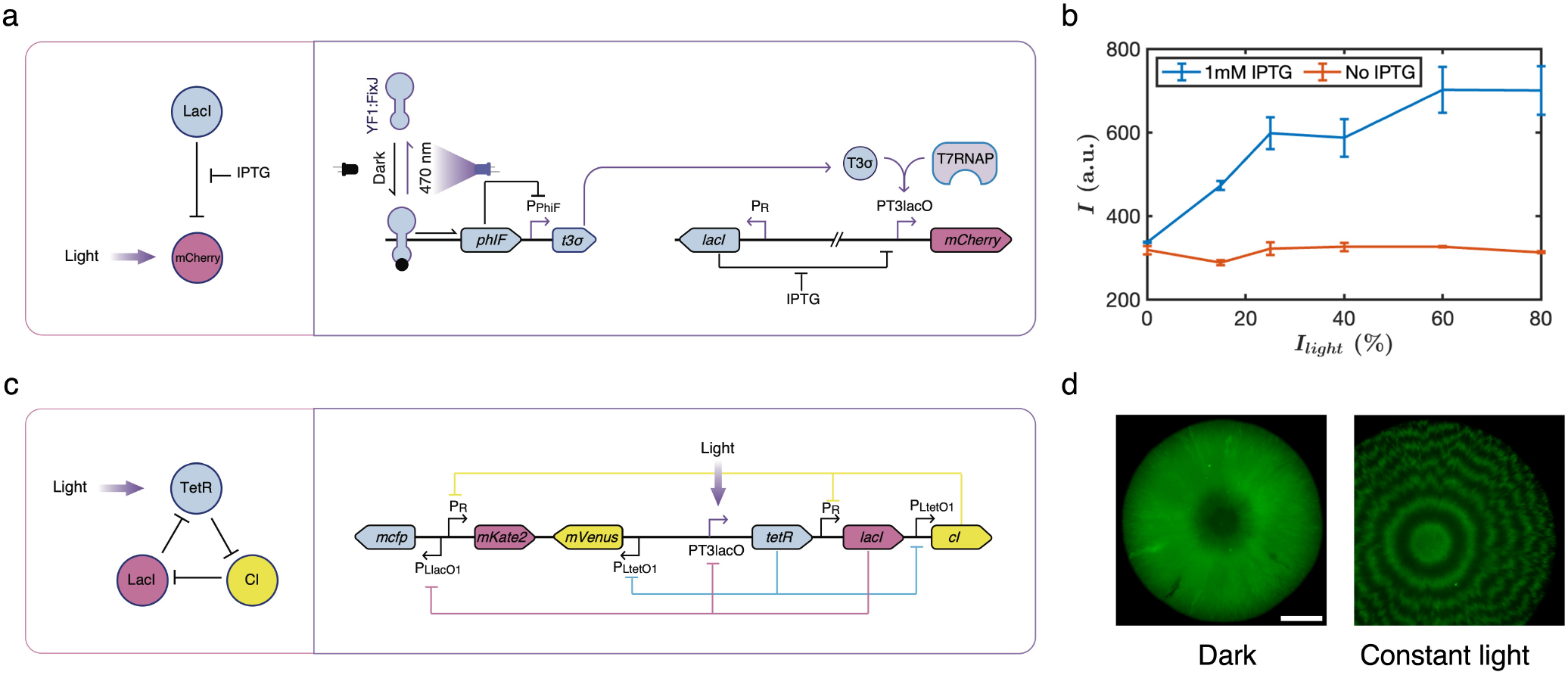
Design and characterisation of the light-inducible repressilator. a) Schematic representation and molecular implementation of the circuit used to charaterise the PT3lacO hybrid promoter. b) Colonies harboring the circuit shown in a) were grown on agar plates in presence or absence of IPTG (1 mM) at different light intensities. The mCherry fluorescence (I, arbitrary units, a.u.) of the colonies was quantified after 96 h of incubation. LacI tightly represses PT3lacO, as cells harboring the circuit were only responsive to light when IPTG was added. Data show the mean of three biological replicates, with error bars depicting the standard deviation (s.d.). c) Schematic representation and molecular implementation of the light-inducible repressilator. d) Colonies harboring the optoscillator circuit only show oscillations (ring patterns) when cultivated under blue light exposure, scale bar = 1 mm.

Once characterised, we took the improved version of the repressilator [23] and exchanged the promoter controlling TetR (PlacO1) with our hybrid promoter (PT3lacO) (Figure 1c). Note that the corresponding CFP reporter still has the original PlacO1 promoter. Thus, CFP is not induced by light. In this manuscript, we always report the fluorescence of the cI/mVenus node downstream of TetR and we display it as green. We first checked whether *E. coli* cells harboring the optoscillator would oscillate when exposed to blue light, but not oscillate in the dark. Indeed, when the plate was covered with aluminum foil for the whole duration of the experiment, the colonies did not display any fluorescent rings, while when exposed to blue light (30 %), rings appeared around a fluorescent centre (Figure 1d), similar to the pattern formed by the regular repressilator [23]. The fluorescent centre can be explained by our experimental setup, where we incubated the colonies the first 20 h in the dark, leading to low TetR and high cI/mVenus expression, before we started the light exposure.

### Synchronization

With continuous light exposure, we noticed that ring patterns were irregular. The irregularity arose from small differences in the period of the individual cells caused by stochastic effects [23]. Over generations these differences increased, meaning that synchronization between the cells decreased with time [38]. As a consequence of this increasing phase difference, the variation in time and position of the next peak increased as the colony grew, resulting in deviations from the initially circular ring pattern. This desynchronisation decreased the average fluorescence amplitude of the rings (Figure 2b).

**Figure 2:**
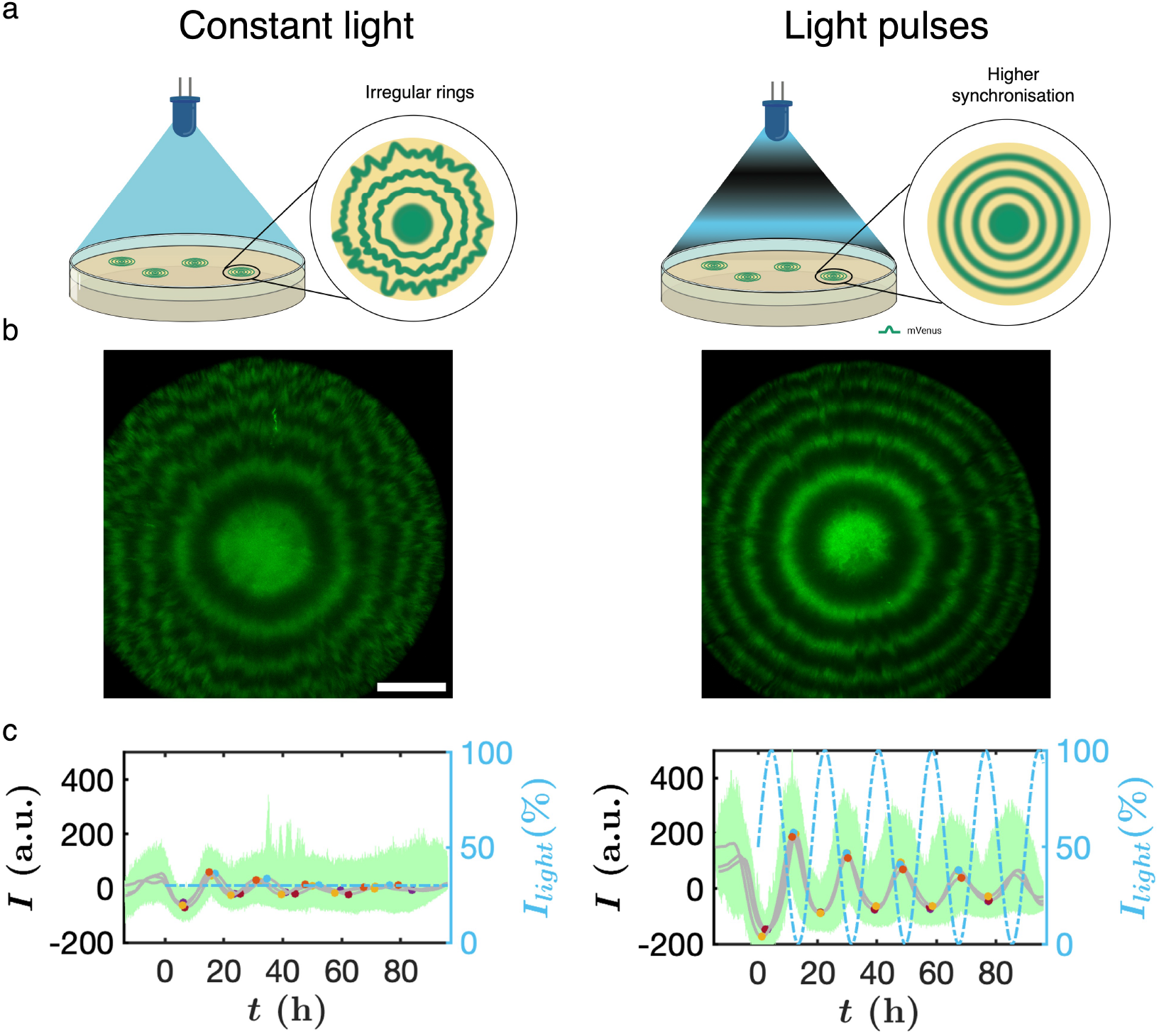
Synchronisation through periodic forcing. a) Representation of the experimental setup for constant light or light pulses with colonies growing on agar plates. b) Picture of colonies grown under constant light (left) and light pulses (right). Better defined rings appear with higher synchronisation of cells under light pulses condition (*T*_*light*_ = 18 h), compared to constant light (30 %); scale bar = 1 mm. c) Each data point of the fluorescence (green) was measured from the center to the border of the colonies and converted to time (details in supplementary information section ). Average fluorescence intensity (*I*) measured for three colonies (grey). The coloured dots show the maxima and minima of oscillations from averaged fluorescence of individual colonies. Light intensity was constant or in a sinusoidal wave shape (dashed blue line).

A simple way to synchronise autonomous self-sustained oscillators is to apply periodic forcing. The individual oscillating cells should re-adjust their phase to the external forcing at each cycle and thus overcome the stochastic effects. To test if our synthetic oscillator could be synchronised by light, we performed experiments with light pulses in a sinusoidal wave shape. Here, the light pulse period is defined as the duration (in hours) required for a full light/dark cycle, with a duty cycle of 50 %. We set the time period of light pulses (*T*_*light*_) to 18 h. After 4 days of growth, colonies showed rings that were sharper and stronger in fluorescence compared to the colonies grown under constant light (Figure 2c). Thus, the synchronisation of the oscillations of the individual cells by the light prevented the amplitude from decreasing and led to sharper and more circular rings in the colonies.

### Resonance

Next, we wondered what would happen if we changed the frequency of light pulses. We expected to find the resonance phenomenon, which occurs when the natural frequency of the driven oscillator (*T*_*osc*_) matches with the frequency of the external forcing. We predicted that this would result in rings with higher fluorescence intensity compared to colonies exposed to different frequencies of periodic forcing. The result from the synchronization experiment already showed strong fluorescent rings, meaning that *T*_*light*_ = 18 h was close to the resonance frequency. Therefore, we repeated the experiment with more *T*_*light*_ conditions, ranging from 12 h to 36 h. We observed the rings with the highest fluorescence intensity – thus resonance – at *T*_*light*_ = 20 h (Figure 3a). In this condition, the rings had the opposite phase with respect to the light pulses. This makes sense as cI and mVenus are inhibited by the light-inducible expression of TetR. Thus, mVenus had the highest expression when the light intensity was the lowest. A more quantitative way to look for resonance is by averaging the amplitude of all rings in a colony and plotting this average fluorescence against the period of the external light pulses. The resonance can be found where the average amplitude is the highest, characterised by a peak in the graph (Figure 3d). As expected we found a resonance peak at 20 h. We also observed another phenomenon typical of forced oscillators: the bacterial oscillations were entrained according to the external light signal. Entrainment was found between *T*_*light*_ = 16 h and *T*_*light*_ = 24 h (Figure 3c). In this interval, the oscillation period increased proportionally with *T*_*light*_ by advancing or delaying the formation of the rings. Above 24 h, the oscillation period is half the period of the light source and we can observe that the oscillator follows the external forcing with every other peak.

**Figure 3:**
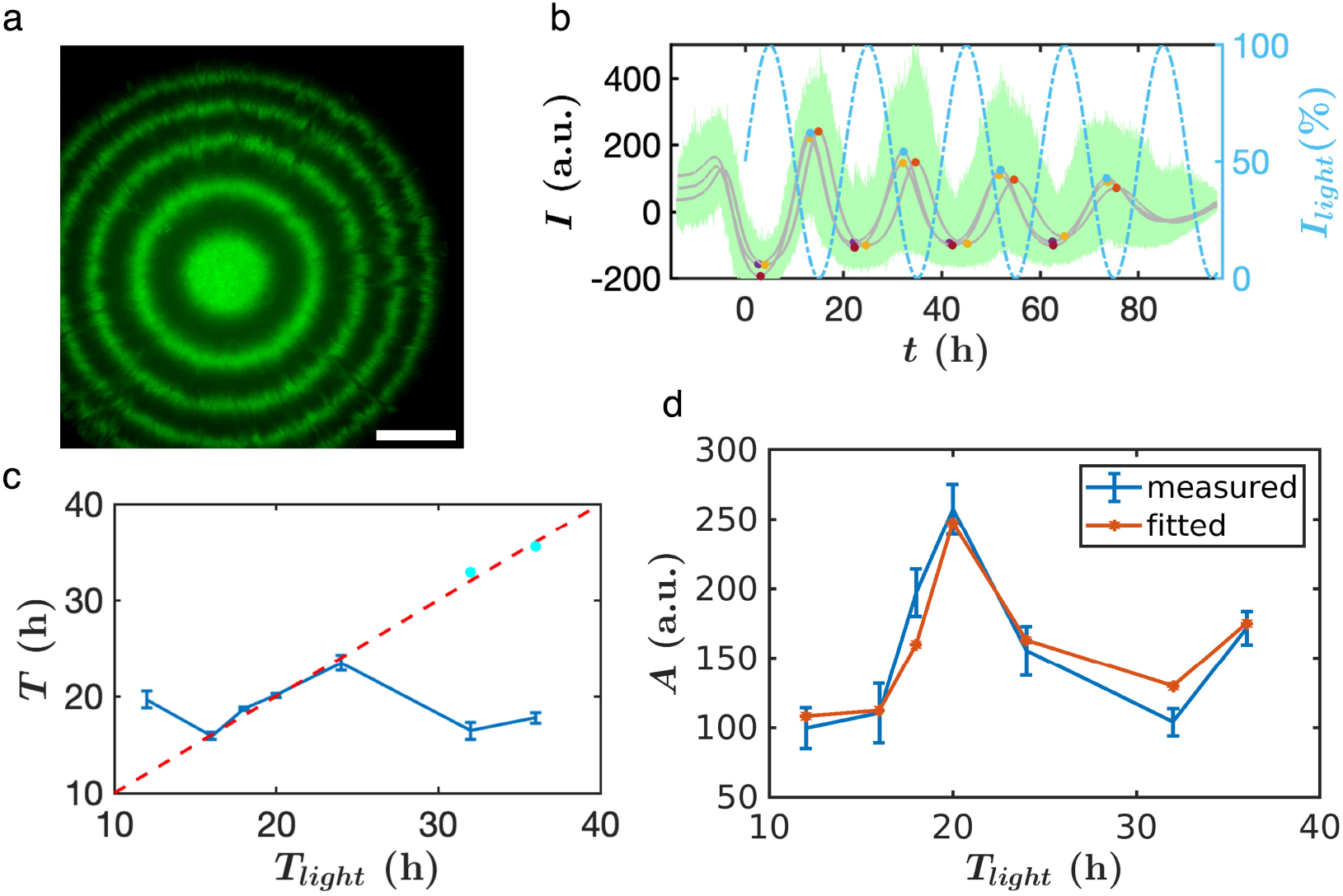
Periodic forcing matching the natural frequency of oscillations leads to resonance. a) Colony showing strong YFP rings (in green) due to the resonance phenomenon, using light pulses at *T*_*light*_ = 20 h. b) Each data point of the fluorescence (green) was measured from the center to the border of the colonies and converted to time (details in supplementary information section ). Average fluorescence intensity (*I*) measured for three colonies (grey). The coloured dots show the maxima and minima of oscillations from averaged fluorescence of individual colonies. Light intensity varied in a sinusoidal wave shape with *T*_*light*_ = 20 h (dashed blue line). c) Evidence of entrainment: the time period of the forced oscillator increases proportionally with the time period of the external forcing from *T*_*light*_ = 16 h to 24 h (dashed red).Above *T*_*light*_ = 24 h, every other oscillation synchronises with the light pulses. We can better visualise this by multiplying the oscillator period by 2 (cyan dots) and see that it falls again in the entrainment region. Data show the mean of three biological replicates, with error bars depicting the s.d. d) The amplitude (A) is the average fluorescence of three replicate colonies for each time period (*T*_*light*_) of the external forcing (blue line) and error bars depict the s.d. The fitted curve from the model is indicated in orange.

### Mathematical model

These interesting observations encouraged us to develop a mathematical model of our genetic circuit to explore whether we might be able to observe other phenomena commonly found in forced oscillators, such as regimes of period doubling or chaos. To do so, we adapted the original mathematical model of the repressilator [22] to represent our optoscillator. Briefly, the system was described with three mRNA species (corresponding to each node) and their respective protein transcription factors. The inhibition of the transcription factors and the light activation was taken into account with Langmuir–Hill functions. The details of the model and the simulation methods can be found in the SI section 2.

We parameterised the model by fitting the data from our hybrid promoter characterisation (Figure 1b) and the data obtained from the resonance experiment (Figure 3d, SI section 2.2). We then used the model to investigate the dynamics of our forced oscillatory system. The deterministic reaction kinetics model showed that – at the resonance frequency – the amplitude of oscillations is significantly larger than in the constant light regime (Figure 4cd). However, the deterministic model cannot describe the details of the desynchronisation, which is an important property in our experiments. Therefore, we performed stochastic reaction kinetics simulations (see the details in SI section 2.2.3) to capture this phenomenon. In Figure 4cIV the red curve shows the average of 1000 independent stochastic simulations. The phase differences between the oscillations of the individual cells increased over time leading to desynchronisation. Due to this stochastic effect, the average amplitude decreased, closely matching our experimental results. On the other hand, in the resonating condition, 1000 independent oscillators remained synchronized and the amplitude did not decrease with time (Figure 4dIV).

**Figure 4:**
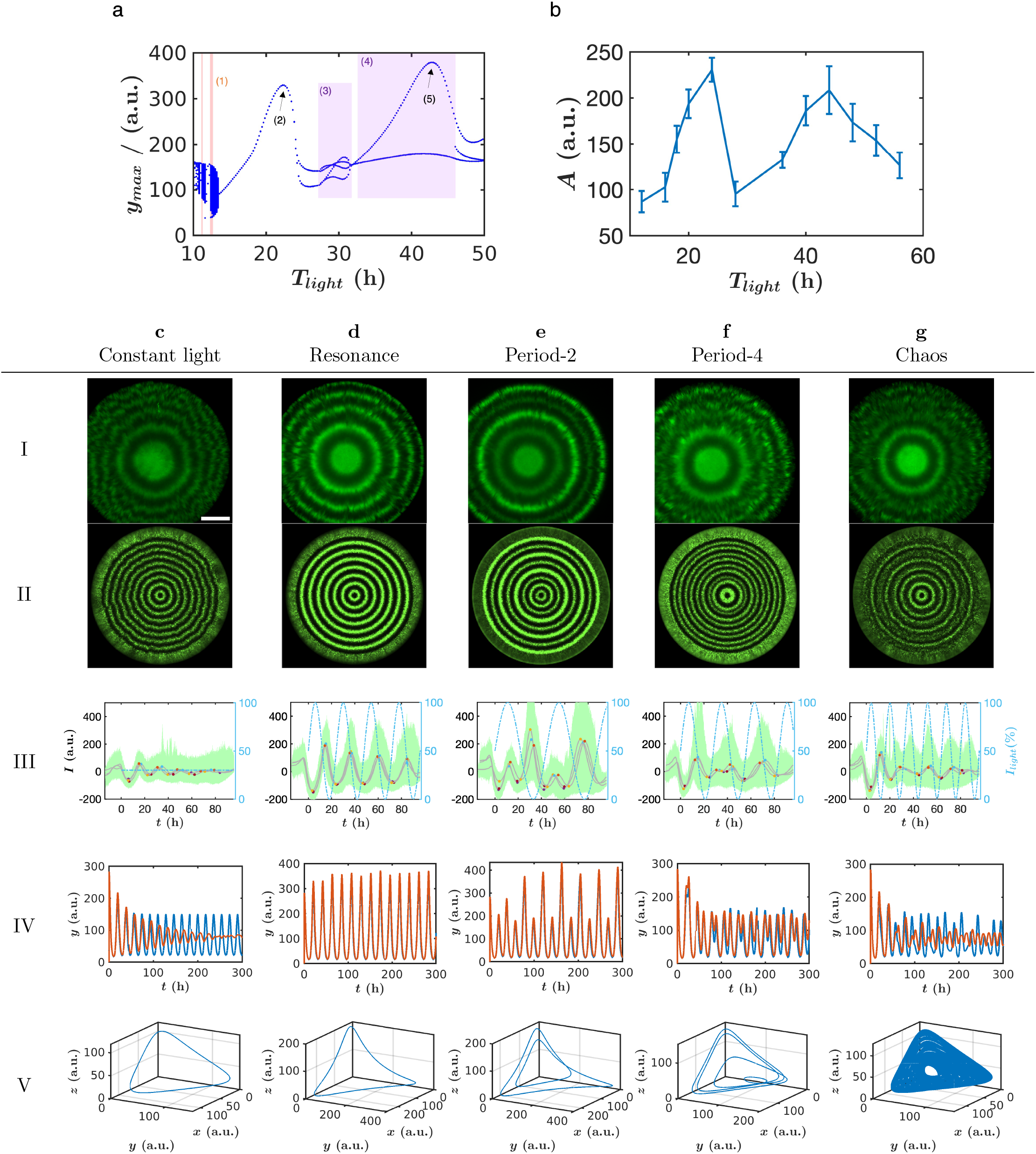
Resonance, undertone, period-2, period-4 and chaos in ring patterns. a) Bifurcation diagram from the model, predicting regions of chaos (1), resonance (2), period-4 (3), period-2 (4) and undertone (5). The average fluorescence (y axis) versus different period forcing (*T*_*light*_). b) Experimental data showing the fluorescence amplitude (A) averaged from three colonies, the error bars are the s.d. It shows the characteristic undertone peak with half the resonance period. c-g) The different regimes: fast desynchronisation with constant light (30 %), resonance at *T*_*light*_ = 24 h, period-2 at *T*_*light*_ = 44 h, period-4 at *T*_*light*_ = 28 h and chaos at *T*_*light*_ = 16 h, respectively. I) Fluorescent picture of a representative colony, scale bar = 1 mm. II) two dimensional reaction–diffusion simulations. III) measured fluorescence from colonies (three replicates per regime in grey), individual fluorescence data points in green, maxima and minima of oscillations from averaged fluorescence of individual colonies as coloured dots and light intensity in dashed blue. IV) deterministic (blue) and stochastic (red) reaction kinetics simulations and V) simulated trajectories in the phase space of the oscillator in different regimes. Further simulation details and results (e.g. Fourier spectra, maxima return map and stroboscopic map) can be found in Figure S5.

To be able to even better compare the simulations to the experimentally observed colonies with rings, we also simulated how the oscillations led to the formation of rings in growing colonies. We carried out two-dimensional reaction–diffusion simulations. The growth of the colony was described with the Fisher–KPP (Kolmogorov—Petrovsky—Piskunov) equation [46], while the cells contained the same stochastic reaction kinetics system that we have described before. The details for the spatial simulations can be found in SI section 2.3. The ring patterns generated from simulations (Figure 4c-gII) were in a good agreement with the experiments (Figure 4c-gI), supporting the validity of our numerical model. We also used the model to examine the effect of the noise in the system (Figure S4 and Supplementary movies). Consistent with the experiment, noise contributes to the irregular shape of the ring patterns under constant light (Figure 4 a), while the resonance regime (Figure 4 b) led to sharper and more circular rings.

Next, we calculated the bifurcation diagram (Figure 4a) as a function of the period of the light pulses to examine whether our optoscillator would show any other interesting dynamical behaviour. The experimental relevant frequency range of the external forcing is limited: if the light source frequency is too high, the cells will not be able to follow it, resulting in the same effect as a constant light regime. Contrarily, if the frequency of the external forcing is too low, we would not get enough rings in a colony to confirm any phenomenon. We therefore limited the bifurcation diagram from 10 h to 50 h. The calculated bifurcation diagram predicted that in addition to resonance, we might be able to observe undertone, period doubling and even chaos (Figure 4a). An undertone is a resonance frequency which occurs at (integral) sub-multiples of the fundamental frequency of the natural oscillator. Our model predicted that we should find another maximum in amplitude at half of the frequency (i.e. at the double period) of the optoscillator. Another interesting phenomenon predicted by our model is the period–doubling bifurcation. In a period-2 oscillation the same peak is repeated every second oscillation, so we can observe bigger and smaller peaks alternately. After two period–doubling bifurcations we can get period-4 (every fourth peak will be repeated) and with an infinitely period–doubling cascade we can also observe chaos. In a chaotic regime the system is deterministic, however, the oscillations are irregular, i.e. the amplitude and the time period are changing and the system is unpredictable [47].

### Undertone, period-2, period-4 and chaos

We thus set out to determine whether we could also observe those additional non-linear phenomena with our optogenetic oscillator. Our model predicted that the amplitude should have a second peak (undertone) located at half of the natural frequency of the oscillator. We therefore repeated the light pulse experiment and extended the maximum time period range of the light induction from 36 h (Figure 3d) to 56 h. Indeed, we observed a second peak in the amplitude at *T*_*light*_ = 44 h (Figure 4b).

The bifurcation diagram also implied that around half of the fundamental frequency of the oscillator we should be able to observe period-2 oscillations. We thus analysed the images of the colonies across different forcing frequencies (Figure 4). In good agreement with the model, we found that colonies grown in the range of 36 h to 48 h time period for light induction showed period-2 oscillations: We observed rings alternating between high and low fluorescence intensities (Figure 4e). The strong rings had the opposite phase to the light pulse, while the weaker rings always occurred in the same phase as the light period. This clearly demonstrated the destructive interference that light imposed on the cI/mVenus node of the oscillator.

In the model, between the resonance and the undertone, we could observe further period doubling (period-4). On the maxima return map (Figure S5G,d), we clearly see four separated dots corresponding to the four different peaks in the period-4 oscillation. Unfortunately the difference between the four peaks is quite small, which makes it difficult to observe a period-4 oscillation experimentally, especially with a limited number of peaks in the colonies. Nevertheless, we were able to observe atypical patterns in this region (*T*_*light*_ = 28 h). In every set of experiments (Figure 4d) we detected many rings of low intensity or even a lack of signal in certain regions of the colonies. These observations can be explained with our model as well: period-4 oscillations lead to oscillations with low amplitude and high frequency. In the experiments these peaks might merge together due to the stochasticity, resulting in either weak or partially disappearing rings.

Finally, our model also predicted a region – occurring before the first resonance peak – where chaotic oscillations might occur (Figure 4a). In a chaotic system, the Laypunov exponent is positive, meaning, that the distance between close trajectories will increase exponentially in time, which makes the system sensitive for initial conditions and the long term behaviour of the trajectories becomes unpredictable. In our simulations we showed that in certain regimes of periodic forcing the system has a positive Laypunov exponent (Figure 4a), the trajectories never repeat themselves in the phase space (Figure 4gV), and the Fourier spectra is continuous, i.e. it does not contain only discrete peaks. Furthermore the maxima return map and the stroboscopic map are continuous curves (Figure S5e), suggesting that our system acts as a chaotic oscillator in this region.

Due to the stochastic nature of the system, there are some variances in the initial state of the cells and, in a chaotic oscillator, these differences will increase in time resulting in desynchronisation resembling the case of oscillations in constant light regime. In the simulations this resulted in irregular and blurred rings (Figure 4gII), while the average amplitude decreased (Figure 4gIV). Some rings were partially disappearing and merging together as the system became progressively more stochastic (from top to bottom in Figure S4 A → B). Indeed in the experiments, colonies grown at *T*_*light*_ = 16 h (Figure 4gI) also displayed irregular and very blurred rings, even more than in the constant light regime. The rapid loss of synchronization is not due to the inability of cells to follow such high frequency of the external forcing, as we used light pulses with square wave shapes at even higher frequency (e.g. T_l_*ight* = 12h) and we still could observe well-defined ring patterns (Figure S1). Overall, the experimental data was surprisingly similar to the simulations, suggesting that our experimental data is consistent with oscillations in a chaotic regime.

## Discussion

Oscillators have been thoroughly studied in every field of natural sciences, but still little is known about their non-linear dynamical properties in biological systems. A well known example – for dumped and forced nonlinear oscillators – is the Duffing equation. In spite of its simplicity, this model can show resonance, and many other interesting dynamical phenomena, such as period doubling bifurcations, chaos and hysteresis [48, 49]. In this work, we were inspired by this approach and constructed a forced synthetic repressilator in *E. coli*. Using a light-inducible system to connect the external forcing to the oscillator, we overcame the need for microfluidics. Through a combination of analysing ring patterns in colonies and mathematical modelling, we have shown that this simple oscillatory circuit can generate complex dynamics depending on the external periodic forcing. In particular, we observed synchronisation, resonance, period doubling (period-2 and period-4) and even chaos.

Our optoscillator resonated when light was shone in sinusoidal waves with *T*_*light*_ between 20 h and 24 h. As the driving period equaled the natural period, bacteria displayed higher synchronisation, higher fluorescence and better defined ring patterns compared to other light regimes. Cells also showed higher synchronisation rate at half of the resonance frequency (undertone), at *T*_*light*_ of 44 h. In fact, these parameter regions where synchronisation is enhanced are called Arnold’s tongues. Apart from resonance, this region allows entrainment of the oscillator to the driving force, a phenomenon observed both in our experiments (Figure 3c) and in prior studies involving synthetic oscillators [37, 38]. It is therefore not surprising to find that many biological oscillators are influenced by periodic environmental cues to keep their physiology and behaviour synchronised [50, 51].

Our experimental data also showed robust period-doubling (period-2 and period-4). To best of our knowledge, no other study presented a synthetic oscillator capable of transforming such dynamic phenomena into spatial patterns. Interestingly, spatial period-doublings have been used to describe wrinkle formation [52, 53] [54] occurring in skin folding [55], mucous membranes [56], the convoluted shape of the brain [57, 58] and biofilm formation [59, 60]. Our circuit could thus be used to generate differential stress conditions in a cell layer in order to reproduce and control wrinkle patterns. This approach could also be interesting to create engineered living materials with desired properties. In fact, controlling wrinkle structure generates surfaces with enhanced properties in soft materials [61, 62].

The combination of our experiments and the mathematical model provides evidence for chaotic oscillations. As far as we are aware, this is the first experimental evidence of a synthetic oscillator producing chaos. For future work, it would be interesting to observe more periods by growing colonies to a bigger diameter with more rings [63] or run the optoscillator in a turbidostat. This would allow us to further characterise this regime. Our finding is consistent with a growing body of literature that points to the importance of chaos in biology [10, 14, 15, 64]. For example, it has been reported that chaos can increase the heterogeneity within a population of genetically identical cells. According to a mathematical model describing the expression of Nuclear Factor-*κ*B (NF-*κ*B), an important transcription factor for the eukaryotic immune response, chaotic oscillations lead to a broader range of expression levels across genes that are regulated by this transcription factor [65]. This heterogeneous gene expression could be beneficial in multi-toxic environments.

As we moved further away from the optoscillator’s natural frequency, synchronization decreased to the point where cells became unable to produce well-defined rings. In the chaotic and period-4 regimes, this desynchonisation effect developed even faster than with constant light. It is therefore a remarkable property of the optogenetic repressilator that depending on the external forcing frequency, the light can either synchronize or desynchronize the oscillations of the individual bacteria. In our simulations at extremely high noise levels in the chaotic regime, the rings became regular again, suggesting an interesting way for stochasticity to control chaos and pattern formation (Figure S4Ee). This stochastic effect on the switch from chaotic to non-chaotic regimes and vice-versa has been previously reported, for example in epidemiological models [66–68].

Our synthetic optoscillator can mimic oscillatory phenomena found in natural systems and help us to distil their underlying design principles. The same optogenetic approach could also be extended to other synthetic oscillators [24, 25, 35, 36], helping us to understand how different topologies respond to external forces. Furthermore, multiple aspects of biological systems are responsive to a varying concentration of inducers in dynamic environments [69, 70]. Consequently, there is a need to explore the behaviour of oscillatory circuits within such scenarios in future research. For example, exploring the behaviour of the optoscillator in gradients of light intensity across space will expand our capacity to manipulate and generate more complex patterns. The ability to control oscillations in gene expression and pattern formation is interesting for a wide range of applications in fields as diverse as biotechnology, medicine, biological sensors, engineered living materials and biocomputing [71–73]. We thus hope our work paves the way for synthetic biology to exploit the rich non-linear dynamics accessible to the optoscillator.

## Methods

### Strains and construction of plasmids

Restriction enzymes were purchased from NEB. DNA fragments were PCR amplified using a high-fidelity DNA polymerase (2x Phanta Max Master Mix, Vazyme). Desalted primers were purchased from Sigma-Aldrich. The DNA fragments were purified using the Monarch PCR & DNA Cleanup Kit (NEB). We assembled the plasmids through Gibson Assembly using the NEBuilder HiFi DNA Assembly Master Mix (NEB) and followed the manufacturer’s protocol. Electrocompetent cells were transformed using electroporation cuvette (2mm gap, Bio-rad) and an electroporator set to 2500 V (Eporator, Eppendorf). The plasmids used in this study are described in Table 1 and primers for PCRs are listed in Table 2.

**Table 1:** Plasmids.

| Name | Function | Reference |
| --- | --- | --- |
| pJFR1 | Template to amplify *t7pol* and *yf1-fixJ* | [39] |
| pJFR2 | Backbone to construct pJP_Azul01 | [39] |
| pJP_Azul01 | Blue light system | This work |
| pLPT41 | Sponge (repressilator) | [23] |
| pLPT107 | Improved repressilator | [23] |
| pJ2072 | Spacer and J23100 for pJP_Ctrl05 | [25] |
| pJP_Ctrl05 | Intermediate for pJP_Ctrl10 | This work |
| pCDF-mCherry | mCherry fragment for pJP_Ctrl05.2 | Our lab |
| pJP_Ctrl05.2 | Intermediate for pJP_Ctrl10 | This work |
| pJP_Ctrl10 | To characterise the hybrid promoter | This work |
| pJP_Rep00 | Light-inducible repressilator | This work |

**Table 2:**
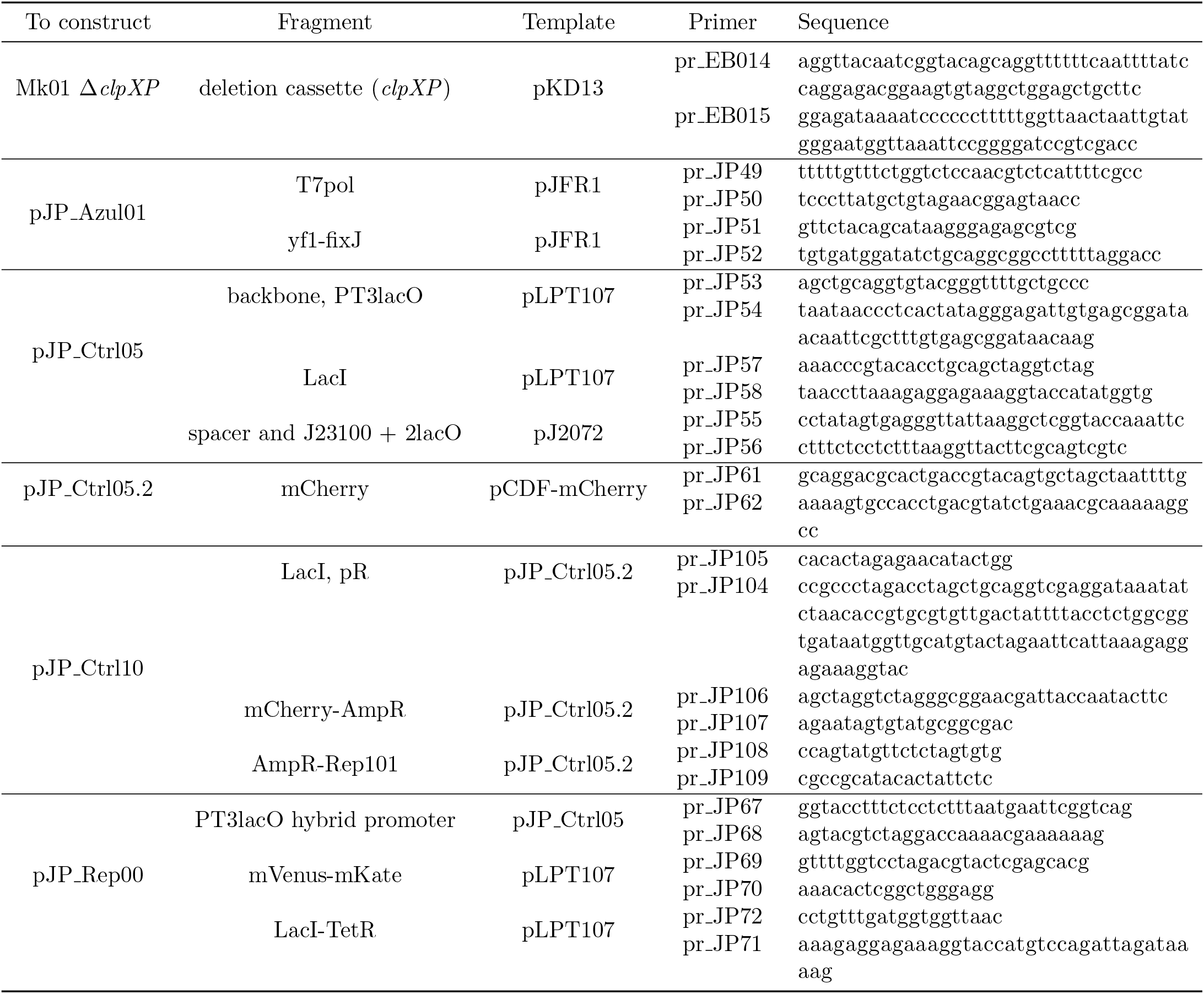
Primers list

To generate the bacterial chassis, we deleted the *clxCP* gene from the *E. coli* MK01 strain [74], which was a kind gift of Sander Tans (Addgene #195090). The genomic deletion was achieved by following the *λ*Red recombination protocol [75]. The DNA fragment for the recombination was generated by PCR amplification of the pKD13 plasmid [75] using primers pr EB14&15.

The light system encoded on the plasmids pJFR1 and pJFR2 [39] was a kind gift of Christopher Voigt. We modified the blue light system from Fernandez-Rodriguez *et al*. [39] to fit it on a single plasmid (pJP Azul01). Briefly, we digested pJFR2 with EcoRI-HF and BsaI-HF v2, while *yf1-fixJ* and *t7pol* were amplified from pJFR1 (using primers pr JP49&50 and pr JP51&52, respectively). After DNA purification, the resulting fragments were assembled together using Gibson assembly and transformed via electroporation into competent *E. coli* NEB5*α*.

The hybrid promoter (PT3lacO) was designed by adding two lac binding sites (lacO) immediately downstream of the PT3 promoter sequence and inserted into pJP Ctrl05. Briefly, we constructed pJP Ctrl05 by amplifying the backbone and *lacI* from pLPT107 using the primers pr JP53&54 (pr JP54 contains the sequence for PT3lacO) and pr JP57&58, respectively, and a spacer fragment from one of our plasmids pJ2072 [25] with the primers pr JP55&56. To avoid any problems related to photo-bleaching with blue light, we changed the fluorescence reporter of the pJP Ctrl05 from mCerulean to mCherry, generating pJP Ctrl5.2. To do so, we digested pJP Ctrl05 with AatII and EcoRI-HF, amplified the mCherry fragment from pCDF-mCherry (from our lab) using primers pr JP61&62 and assembled the two sequences by Gibson assembly. To more accurately represent the TetR node of the repressilator the J23100 promoter controlling the *lacI* transcription was subsequently replaced by the pR promoter. This was done by PCR amplification of pJP Ctrl5.2 using the primers pr JP104&105, pr JP106&107 and pr JP108&109, followed by Gibson assembly. The final version of this plasmid, which was used for the characterisation of the PT3lacO promoter, is called pJP Ctrl10.

For the light-inducible repressilator (pJP Rep00), we started with plasmids pLPT107 and pLPT41 [23], which were kind gifts from Johan Paulsson (Addgene plasmid #85525 and #85524). To construct pJP Rep00, we deleted the promoter pLtetO upstream of *tetR* and replaced it with PT3lacO. To do so, we digested pLPT107 with HpaI and StuI to obtain the backbone and we amplified the remaining fragments from pLPT107 using the primers pr JP69&70 and pr JP71&72, and the hybrid promoter (PT3lacO) from pJP Ctrl05 using primers pr JP67&68.

Plasmids pJP Azul01 and pJP Rep00 will be deposited at Addgene.

### Optogenetic experiments

#### Optogenetic setup

For constant and light pulses conditions, we used a LITOS device ([45]) for light induction. This device is composed of a LED RGB matrix and a control unit to which we can upload a program to set the stimulation time, wavelength (R = 620 - 630 *η*m, G = 520 - 525 *η*m, B = 465 - 470 *η*m) and intensity and select the positions of the LEDs we want to turn on. In this work, we only used the blue LED, therefore the red and green intensities were set to 0.

To convert the relative light intensities (15 %, 20 %, 25 %, 30 %, 35 %, 40 %, 60 %, 80 % and 100 %) to absolute light values, we measured the light emitted by LITOS in a dark room, using the Skye single channel light sensor device (SKYE instruments LTD). The given unit (*×*10*µmol* · *m*^−2^ · *s*^−1^) was then converted to W·m^−2^. The graph correlating the light intensities in percentages to absolute values is shown in Figure S2.

#### Growth of colonies

To test the PT3lacO hybrid promoter, pJP Ctrl10 and pJP Azul01 were co-transformed into *E. coli* MK01 Δ*clxCP*. A colony was inoculated in a tube containing 3 mL of liquid LB with ampicillin (100 *µ*g/ mL) and spectinomycin (50 *µ*g/ mL).

After an overnight incubation at 30 °C, cells were serial diluted 4x10^7^ times. 100 *µ*L were homogeneously spread on LB agar plates (90 mm in diameter) with ampicillin (100 *µ*g/ mL), spectinomycin (50 *µ*g/ mL) and with or without 1 mM IPTG. The plates were wrapped in aluminum foil and incubated at 30 °C for 20 h. Next, the aluminum foil was removed for microscopy imaging to measure the initial size of the colonies. Each plate contained 5 to 10 colonies, 3 (replicates) of them were randomly selected and marked to track initial and final size of respective colonies. Then, the plates were placed upside-down on the LITOS LED matrix with different light conditions (constant light at 15, 20, 25, 30,35, 40, 60 and 80 % of light intensity). The LITOS setup with the plates was placed in a Peltier incubator (IPP 750 Plus, Memmert GmbH). The temperature was set to 21 °C and the experiment ran for 96 h. After that, colonies were imaged with a fluorescence microscope to measure fluorescence and the final size of the colonies (see microscopy section for details).

For the ring patterns experiment, we first obtained the optoscillator strain by co-transforming pLPT41, pLPT107 and pJP Azul01 into *E. coli* MK01 Δ*clxCP*. The experimental procedure was the same as described above, except that IPTG was not added to the medium and the antibiotics used for inoculation were kanamycin (50 *µ*g/ mL), ampicillin (100 *µ*g/ mL) and spectinomycin (50 *µ*g/ mL). For the constant light condition, light intensity was set to 30 % until the end of the experiment, and for the light pulses regimes, the intensity of light (*I*) varied from 0 to 100 % in a sinusoidal wave shape (*I* = *I*_0_(1 + *sin*(2*π/T*_*light*_))*/*2) according to the chosen time period (*T*_*light*_). The intensity was set with an eight bit resolution, and we changed the intensity twenty times during one time period.

#### Microscopy

Pictures were taken with an AxioImager M1 microscope (Zeiss) with an Achromat 2.5X/0.12 Fluar objective and an EMCCD camera (Photometrics) controlled by the VisiView 7.5 software (Visitron Systems).

Filters setup: DsRed and YFP channels were used to measure the fluorescence of mKate2 (and mCherry) and mVenus, respectively. The exposure time of every channel was set to 100 ms, including the brightfield channel.

#### Image analysis

Image processing steps to generate quantitative data from the fluorescent ring patterns were carried out in MATLAB (R2023b). We first measured the fluorescence intensity across the radius of the colony (space vs intensity). To determine the oscillator’s time period and to compare its phase to the light pulse period, we transformed the space vs intensity data from our ring pattern experiments to time vs intensity data. Because the bacterial colony does not expand linearly, we first measured a growth curve correlating the radius of the colony and the elapsed time (space vs time). With that, we could transform the space vs intensity plots to time vs intensity graphs. The details of this transformation are described in the SI section .

### Mathematical model and simulations

The model is described in detail in the supplementary information section 2. Briefly, the deterministic reaction kinetics equations were described with ordinary differential equations (ODEs) and they were solved in a dimensionless form in MATLAB (R2023b) with the Simbiology toolbox, while the stochastic differential equations (SDEs, based on the chemical Langevin equation (CLE)) were solved in C++ with the fully composite Patankar method [76] (SI section 2.3.2).

To compare the experimentally observed patterns with the model, we also performed spatial simulations. Details are described in the supplementary information section 2.3. Briefly, simulations are based on the Fisher–KPP (Kolmogorov— Petrovsky-–Piskunov)[46] equations. These partial differential equations (PDEs) were solved in C++ with the finite difference method, with the forward time–centered space (FTCS) scheme, while the stochastic reaction kinetics equations were solved using the fully composite Patankar method [76] .

## Contributions

J.P, G.H and Y.S designed the experimental research. J.P performed the experiments, G.H performed the mathematical modeling, J.P and G.H analysed the data and prepared the corresponding figures. J.P, G.H and Y.S wrote the manuscript. All authors have given approval to the final version of the manuscript.

## Acknowledgements

We thank Emanuele Boni for deleting *clpXP* from *E. coli* MK01. We also thank Dora Takiya Bonadio for helping us with the schematic images. This work was funded by a Swiss National Science Foundation (310030 200532 awarded to Y.S), a fellowship of the Agassiz foundation (awarded to JP), a UNIL FBM PhD fellowship in Life Sciences (awarded to JP) and the University of Lausanne.

