## Supplementary Information for "From resonance to chaos: modulating spatiotemporal patterns through a synthetic optogenetic oscillator"

### 1 Supplementary experiments

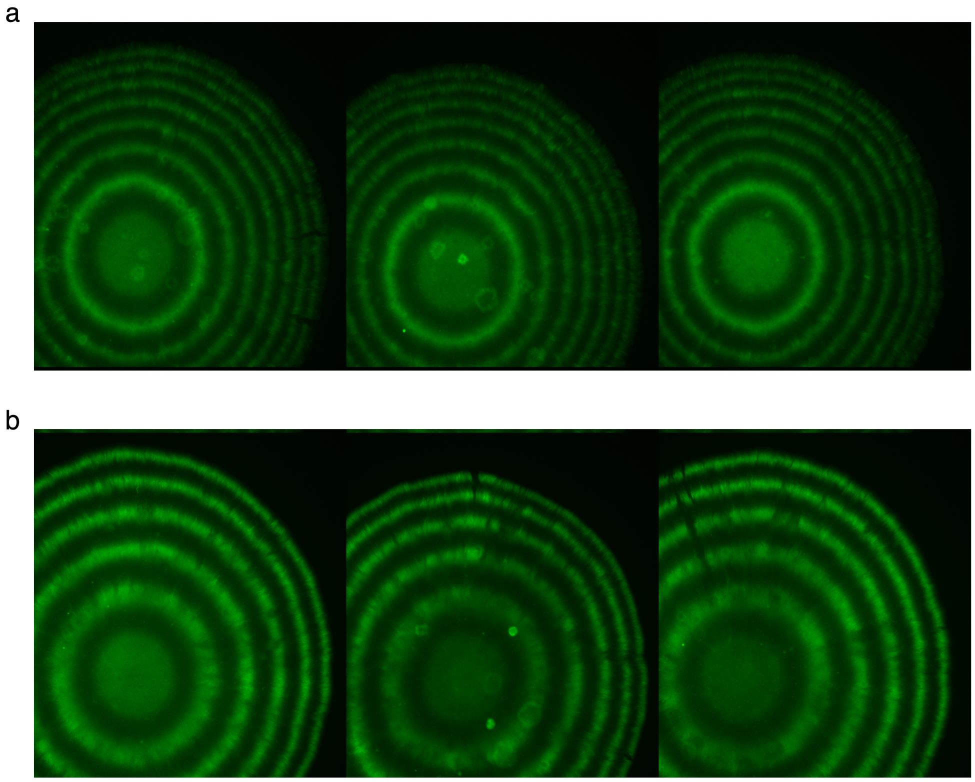


**Figure S1:** Square wave pulses with light intensity varying from 0 to 100%. a) $T_{light} = 12$ h. b) $T_{light} = 18$ h. Colonies harboring the optoscillator were subjected to square wave shape of light induction. The cells were capable of following the light pulses and produce rings with high frequency (e.g. $T_{light} = 12$ h). This demonstrates that the loss of pattern in the chaotic regime ($T_{light} = 16$ h) is not due to cell's inability to follow higher frequency pulses.

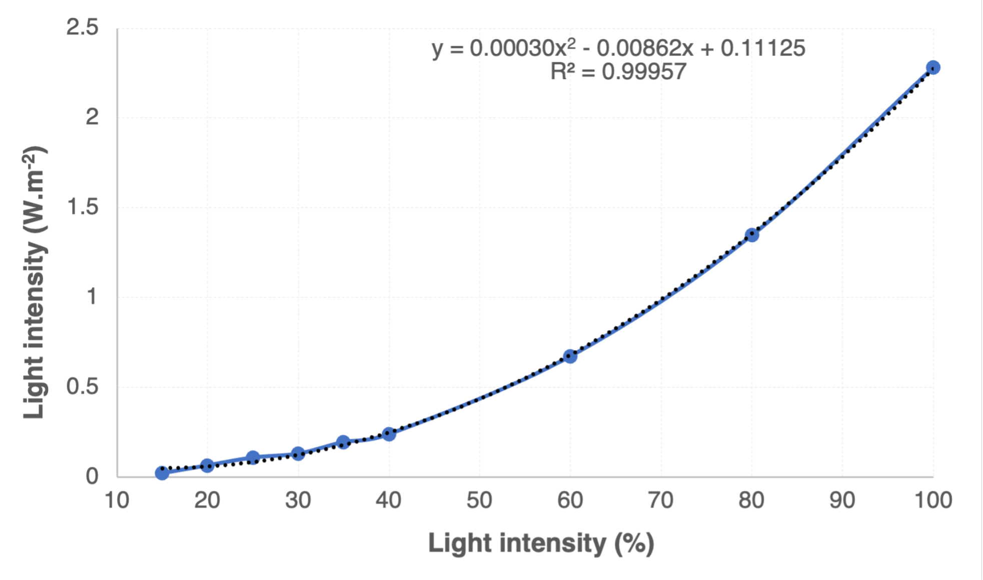


**Figure S2:** Conversion of relative light intensities (%) from the LITOS device absolute values in W$\cdot$m$^{-2}$. The light emitted by LITOS was measured with a light sensor device.

### 2 Model description

#### 2.1 The model

Our model is based on the original model of the repressilator[1]. It contains six species: three mRNAs ($m_x$, $m_y$, $m_z$) and the three corresponding proteins ($P_x$, $P_y$, $P_z$). The model takes into account the degradation rate of the mRNAs and the protein transcription factors ($k_2$ and $k_4$) and as a simplification, the parameters are the same for all mRNAs and proteins. The binding of the transcription factor to the DNA, decreases the mRNA production rate ($k_1$) and the inhibition was described by the Langmuir–Hill function (with the parameters: $K$, $n_P$), while $k_0$ represents the leakage of the promoters (i.e. mRNA production rates). The protein production is proportional to the corresponding mRNA concentration and its rate constant is $k_3$.

The original model was extended with a light–inducible promoter and the light activation was taken into account with a Langmuir–Hill function (Table S1). The maximum strength of the light-inducible promoter (maximum $m_x$ production rate) is $k_{1x}$, the leakage of the light-inducible promoter is $k_{0,x}$, while the half-saturation constant and the Hill exponent are $K_I$ and $n_I$, and $I$ is the relative intensity of the light source (its value is given in % compared to its maximum value).

**Table S1:** The original reaction kinetics model of the repressilator is extended with a light-inducible promoter. The meaning of the parameters can be found in the text.

| Reaction | Rate law |
|---|---|
| $\emptyset \longleftrightarrow \mathrm{m}_x$ | $r_{1,x} = k_0 + \left( k_{0x} + \dfrac{k_{1x}\left(\dfrac{I}{K_I}\right)^{n_I}}{1+\left(\dfrac{I}{K_I}\right)^{n_I}} \right) \dfrac{1}{1+\left(\dfrac{[P_z]}{K}\right)^{n_P}} - k_2[m_x]$ |
| $\emptyset \longleftrightarrow \mathrm{m}_y$ | $r_{1,y} = k_0 + \dfrac{k_1}{1+\left(\dfrac{[P_x]}{K}\right)^{n_P}} - k_2[m_y]$ |
| $\emptyset \longleftrightarrow \mathrm{m}_z$ | $r_{1,z} = k_0 + \dfrac{k_1}{1+\left(\dfrac{[P_y]}{K}\right)^{n_P}} - k_2[m_z]$ |
| $\mathrm{m}_i \longleftrightarrow \mathrm{m}_i + \mathrm{P}_i$ | $r_{2,i} = k_3[m_i] - k_4[P_i], \quad i \in \{x, y, z\}$ |

##### 2.1.1 Creating the dimensionless model

The model contains 11 parameters. To reduce the number of parameters, the following assumptions were made: $k_0 = k_1/10^3$, $n_P = 2$ (based on the original repressilator model[1]), $k_{0x} = k_{1x}/10^2$ (the light-inducible system has a higher leakage [2]), which results in 8 unknown parameters. If we are using different units for the mRNA and the proteins, then there are three dimensions $(M_P, M_m, s)$ and the dimensionless model can be described with only five parameters. After choosing the following units:

$$M_P = [K], \quad s = \left[\frac{1}{k_4}\right], \quad M_m = \left[\frac{k_4 K}{k_3}\right], \tag{1}$$

we get the following dimensionless parameters and concentrations:

$$\hat{k}_1 = \frac{k_1 k_3}{k_4^2 K}, \quad \hat{k}_{1x} = \frac{k_{1x} k_3}{k_4^2 K}, \quad \hat{k}_2 = \frac{k_2}{k_4}, \quad [\hat{m}_i] = \frac{k_3 [m_i]}{k_4 K}, \quad [\hat{P}_i] = \frac{[P_i]}{K}, \tag{2}$$

and the following parameters have unit values: $\hat{k}_3 = \hat{k}_4 = \hat{K} = 1$. The dimensionless model can be found in Table S2.

**Table S2:** The dimensionless model contains the following parameters: $K$, $\hat{k}_1$, $n_I$, $\hat{k}_2$, $\hat{k}_{1x}$. The fitted values of the parameters can be found in Table S4.

| Reaction | Rate law |
|---|---|
| $\emptyset \longleftrightarrow \mathrm{m}_x$ | $\hat{r}_{1,x} = \dfrac{\hat{k}_1}{10^3} + \left( \dfrac{\hat{k}_{1x}}{10^2} + \dfrac{\hat{k}_{1x}\left(\dfrac{I}{K_I}\right)^{n_I}}{1+\left(\dfrac{I}{K_I}\right)^{n_I}} \right) \dfrac{1}{1+\left[\hat{P}_z\right]^2} - \hat{k}_2[\hat{m}_x]$ |
| $\emptyset \longleftrightarrow \mathrm{m}_y$ | $\hat{r}_{1,y} = \dfrac{\hat{k}_1}{10^3} + \dfrac{\hat{k}_1}{1+\left[\hat{P}_x\right]^2} - \hat{k}_2[\hat{m}_y]$ |
| $\emptyset \longleftrightarrow \mathrm{m}_z$ | $\hat{r}_{1,z} = \dfrac{\hat{k}_1}{10^3} + \dfrac{\hat{k}_1}{1+\left[\hat{P}_y\right]^2} - \hat{k}_2[\hat{m}_z]$ |
| $\mathrm{m}_i \longleftrightarrow \mathrm{m}_i + \mathrm{P}_i$ | $\hat{r}_{2,i} = [\hat{m}_i] - \left[\hat{P}_i\right], \quad i \in \{x, y, z\}$ |

**Stochastic simulations** In the stochastic reaction kinetics simulations the chemical Langevin equation (CLE)[3, 4] was used:

$$\frac{\mathrm{d}\vec{c}}{\mathrm{d}t} = \sum_{i=1}^{M} \nu_i r_i(\vec{c}, t) + \frac{1}{\sqrt{\Omega}} \sum_{i=1}^{M} \nu_i \sqrt{r_i(\vec{c}, t)}\ \Gamma_i(t), \tag{3}$$

where the sum goes for every reaction, $\nu_i$ is the stoichiometry coefficient, $\Gamma_i(t)$ is the Wiener process (uncorrelated independent Gaussian white noise, $\Gamma_i = \lim_{dt \to 0+} \frac{\mathcal{N}(0,1)}{\sqrt{dt}}$), $\Omega$ is the volume of the cell, and $\vec{c}$, $t$ and $r_i$ are the concentration vector, time and the reaction rate of the $i$th reaction respectively.

#### 2.2 Fitting the parameters

##### 2.2.1 The Langmuir–Hill coefficients

To calculate the Langmuir–Hill coefficients, we designed and characterized a two-node light-inducible circuit (Figure S3/A, Figure 1/ab). The light induces the expression of mCherry. At the same time, mCherry expression can also be repressed by the constitutively expressed repressor LacI and the repression of LacI can be reduced by addition of IPTG to the medium.

Figure S3 shows how the mCherry fluorescence of the colonies is increasing with the intensity of the light source due to the activation of the light-inducible promoter in the presence of 1 mM IPTG.

The measured fluorescence signal($I_R$) is proportional to the concentration of the fluorescent mCherry protein ($[\hat{P}_x]$):

$$I_R = a + b[\hat{P}_x]. \tag{4}$$

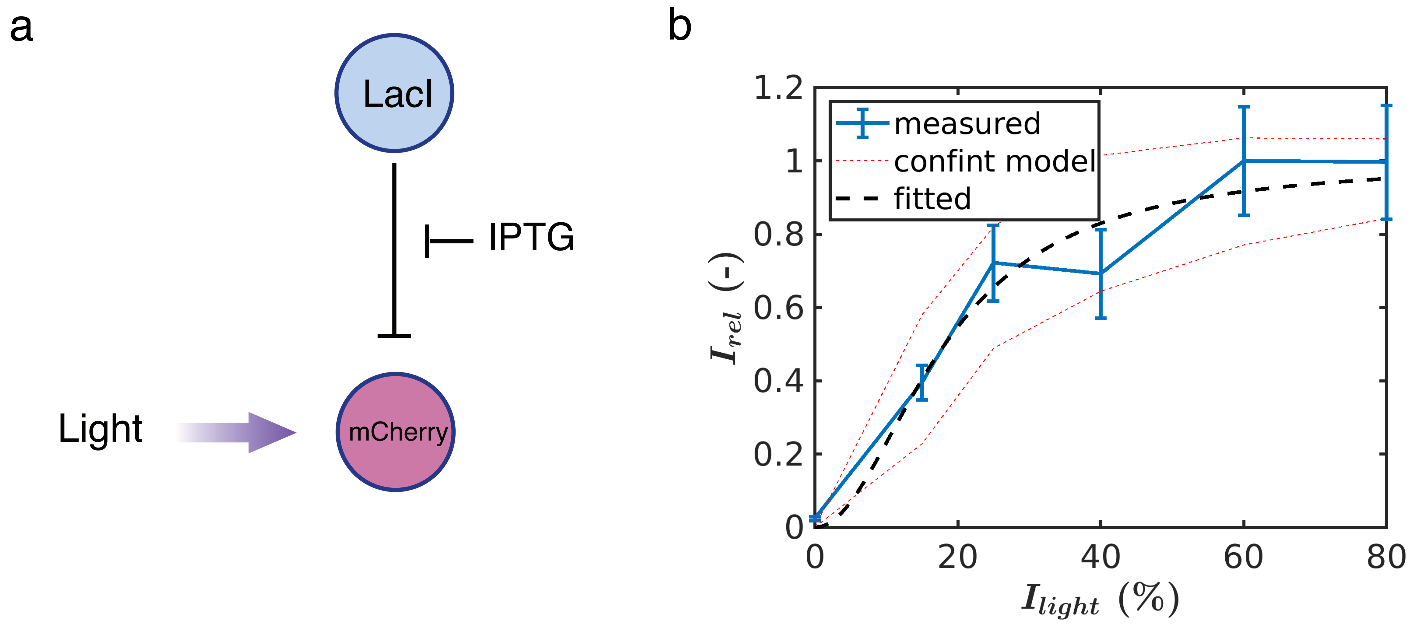


**Figure S3:** a) The two-node light-inducible circuit, b) The relative fluorescence $I_{rel}$ as a function of the external light intensity $I_{light}$ in the presence of high IPTG concentration (1 mM). The fluorescence of the colonies were determined from the same radius range for every colony (from 100 to 300 pixel). The minimum fluorescence (from a colony without IPTG in the dark) was subtracted from the signal and it was normalized with the maximum fluorescent signal (80 % light, high IPTG concentration). The experimental data show the mean of three biological replicates (blue line), with error bars depicting the s.d. of the normalized replicates. For the model, the fitted Hill–function is shown in dashed black line, with the 95 % confidence interval (dashed red).

**Table S3:** The simplified dimensionless model based on Table S2 for the two-node system shown in Figure S3. $\hat{r}_{1,x}$ is the rate law for $m_x$ production in the absence of IPTG, while $\hat{r}_{1,x}^{\mathrm{IPTG}}$ denotes the rate equation in the presence of a high concentration of IPTG.

| Reaction | Rate law |
|---|---|
| $\emptyset \longleftrightarrow \mathrm{m}_x$ | $\hat{r}_{1,x} = \dfrac{\hat{k}_1}{10^3} + \left( \dfrac{\hat{k}_{1x}}{10^2} + \dfrac{\hat{k}_{1x}\left(\dfrac{I}{K_I}\right)^{n_I}}{1+\left(\dfrac{I}{K_I}\right)^{n_I}} \right) \dfrac{1}{1+\left[\hat{P}_z\right]^2} - \hat{k}_2[\hat{m}_x]$ |
| | $\hat{r}_{1,x}^{\mathrm{IPTG}} = \dfrac{\hat{k}_1}{10^3} + \dfrac{\hat{k}_{1x}}{10^2} + \dfrac{\hat{k}_{1x}\left(\dfrac{I}{K_I}\right)^{n_I}}{1+\left(\dfrac{I}{K_I}\right)^{n_I}} - \hat{k}_2[\hat{m}_x]$ |
| $\emptyset \longleftrightarrow \mathrm{m}_z$ | $\hat{r}_{1,z} = \dfrac{\hat{k}_1}{10^3} + \hat{k}_1 - \hat{k}_2[\hat{m}_z]$ |
| $\mathrm{m}_i \longleftrightarrow \mathrm{m}_i + \mathrm{P}_i$ | $\hat{r}_{2,i} = [\hat{m}_i] - \left[\hat{P}_i\right], \quad i \in \{x, z\}$ |

Without the $y$ node (omitting $m_y$ and $P_y$ from the original model) we get the model of the two-node subsystem shown in Figure S3a (Table S3). If the colony is growing for a long time without changing the experimental conditions, the protein and mRNA concentrations will be close to the fix point of this system. From $\hat{r}_{1,z}$ and $\hat{r}_{2,z}$ the fix point of $[\hat{m}_z]$ and $[\hat{P}_z]$ are: $[\hat{P}_z]^\star = [m_z]^\star = 1.001 \cdot \hat{k}_1/\hat{k}_2$ and similarly $[\hat{P}_x]^\star = [m_x]^\star$ so the fix point for the fluorescence can be calculated based on Eq. 4 and $\hat{r}_{1,x}$:

$$I_R^\star = c_1 + \left( c_2 + c_3 \frac{\left(\frac{I}{K_I}\right)^{n_I}}{1 + \left(\frac{I}{K_I}\right)^{n_I}} \right) \frac{1}{1 + \left(\frac{[\hat{P}_z]^\star}{K}\right)^{n_P}}, \tag{5}$$

where $c_1 = a + \frac{b\hat{k}_1}{10^3 \hat{k}_2}$, $c_2 = \frac{b\hat{k}_{1x}}{10^2 \hat{k}_2}$, $c_3 = \frac{b\hat{k}_{1x}}{\hat{k}_2}$, while in the presence of high IPTG concentration (i.e. without the repression of $P_z$):

$$I_{R1}^\star \equiv I_R^\star([IPTG]_{high}, I) \approx c_1 + c_2 + c_3 \frac{\left(\frac{I}{K_I}\right)^{n_I}}{1 + \left(\frac{I}{K_I}\right)^{n_I}}. \tag{6}$$

The maximum fluorescent mCherry signal ($I_{R1}^{max,\star}$) was measured with high light intensity (80 %) and with high IPTG concentration (1 mM). The maximum fluorescence intensity can be calculated from the model:

$$I_{R1}^{max,\star} \equiv I_R^\star([IPTG]_{high}, I_{high}) \approx c_1 + c_2 + c_3. \tag{7}$$

On the other hand, in dark and in the absence of IPTG, the fluorescent signal is minimal, it only contains the leakage and the background fluorescence($I_{R2}^\star$). The calculated minimum fluorescence intensity is:

$$I_{R2}^\star \equiv I_R^\star([IPTG] = 0, I = 0) \approx c_1. \tag{8}$$

We measured the fluorescence as a function of the external light intensity ($I_{R1}^\star$) when the inhibition by LacI was repressed with IPTG. After subtracting the minimum fluorescence ($I_{R2}^\star$) and normalizing it with the maximum fluorescence ($I_{R1}^{max,\star}$) we got the following dimensionless fluorescence, which can be described with a Langmuir–Hill function:

$$\hat{I}_R^\star = \frac{I_{R1}^\star - I_{R2}^\star}{I_{R1}^{max,\star} - I_{R_2}^\star} \approx \frac{\left(\frac{I}{K_I}\right)^{n_I}}{1 + \left(\frac{I}{K_I}\right)^{n_I}}. \tag{9}$$

We used the Levenberg–Marquardt nonlinear least squares algorithm[5] weighted with the reciprocal of the experimental error (nlinfit function of MATLAB, with $10^{-10}$ tolerance on the estimated coefficients and $10^{-14}$ tolerance for the residual sum of squares).

The fitted curve can be found in Figure S3. With this independent experiment on the light system – on a subsystem of the forced oscillator – we have determined two of the five model parameters: $n_I$ and $K_I$. Their values can be found in Table S4.

##### 2.2.2 The resonance curve

To determine the three remaining parameters ($\hat{k}_1$, $\hat{k}_{1x}$, $\hat{k}_2$), we fitted the model to the resonance peak (Figure 3d). In the parameter estimation dimensionless parameters were used to be able to compare the experimental and simulation results. The amplitude $A$ was divided with the average amplitude $< A >$, while the time period ($T_{light}$) of the light source was divided with the time period of the oscillator at constant 30 % light intensity ($T_{30\%}$). Later on these values were used to scale the dimensionless time and amplitude to real scale: $T_{scale} = T_{30\%}^{meas}/T_{30\%}^{sim} = 1.6864$ h, $A_{scale} =< A >_{meas} / < A >_{sim}= 2.4287$ a.u. (where "sim" denotes the simulation, while "meas" denotes the measured values).

To calculate the resonance curve in the simulations, first the time period of the oscillator was determined at 30 % light intensity ($\hat{T}_{30\%}$), then the dimensionless time period of the external light source ($\hat{T}_{light}$) was calculated according to the experimental setup: $\hat{T}_{light} = T_{light}/T_{30\%} \cdot \hat{T}_{30\%}$. After that, the average amplitude ($\hat{A}$) of the oscillation of $P_y$ was calculated according to the external light time periods $\hat{T}_{light}$ and the amplitude values were divided with their average $<\hat{A}>$.

The original parameters of the repressilator model[1] ($p_0$) were used as a starting point in the model fitting and the parameters values were constrained for the $[0.5p_0, 2p_0]$ range to stay in a physically relevant parameter space. We prescribed furthermore that the maximal strength of the light-inducible promoter ($\hat{k}_{1x}$) should be smaller than the other (non light-inducible) promoters ($\hat{k}_1$).

The sum of squared error (weighted with the experimental error) was minimised first with the simplex method[6] (fminsearch function of MATLAB), then the result was fine-tuned with the Levenberg–Marquardt nonlinear least squares algorithm[5] (nlinfit function of MATLAB, with $10^{-8}$ tolerance for the parameters and $10^{-10}$ function tolerance). The fitted values can be found in Table S4.

**Table S4:** The fitted dimensionless model parameters. Every parameter was fitted in a logarithmic form, except the Hill coefficient of the light system ($n_I$). The fitted value is given with the 95 % confidence interval of the fitting.

| Parameter | Value | Parameter | Value | Parameter | Value |
|---|---|---|---|---|---|
| $\log_{10}(K)$ | $1.26 \pm 0.13$ | $\log_{10}(\hat{k}_1)$ | $3.77 \pm 0.36$ | $n_I$ | $2.0 \pm 1.6$ |
| $\log_{10}(\hat{k}_2)$ | $0.10 \pm 0.03$ | $\log_{10}(\hat{k}_{1x})$ | $3.75 \pm 0.89$ | | |

##### 2.2.3 Simulation details

**Deterministic model** The deterministic simulations were performed with the sundials solver in MATLAB with a relative tolerance $10^{-8}$ and an absolute tolerance $10^{-10}$. The experiments started from a dark state, so the initial conditions were determined according to the fixed point of the ODE system with $\hat{I} = 0$ (see Table S5 for further details). For the simulations where we compared the trajectories with the experimental time–intensity curves – and the initial transient state was of interest –, the simulation length was 200. When the initial transient period was not used, the data collection started after a given time ($\hat{T}_0$). In the calculation for the bifurcation diagram (Figure 4) $\hat{T}_0 = 400$ and the simulation time was $\hat{T}_{sim} = 10^3$, while for all the other simulations (calculation of the Fourier spectra, the stroboscopic map, the maxima return map and the phase space) $\hat{T}_0 = 10^3$ and the length of the simulations was $\hat{T}_{sim} = 10^4$.

**Stochastic model** The stochastic differential equations (SDEs) were solved using the fully composite Patankar–Euler method[7]. In this method, the solution is calculated in every time step with the Euler–Maruyama method[8]. However, if the noise or the step size is too large, one can get negative concentration values. In this case the concentration is recalculated with the deterministic Patankar method and if the concentration is still negative, then the stochastic Patankar method is used[7]. We implemented this method in C++ and the solution was parallelised with OpenMP. 1000 simulations were carried out for each case and their average was used for further analysis. The system size was chosen to be $\hat{\Omega} = 5$ and a time step was $\hat{dt} = 2 \cdot 10^{-3}$.

#### 2.3 Spatial simulations

To visualize the patterns that our model predicted, spatial simulations were performed. The growth of the colony was described with the Fisher–KPP (Kolmogorov—Petrovsky-–Piskunov)[9] equation:

$$\frac{\partial c}{\partial t} = D\nabla^2 c + kc(F - c), \tag{10}$$

where $c$ is the concentration of the cells on the surface and $D$, $k$, $F$ are the diffusion coefficient, the rate constant of the logistic growth and the carrying capacity, respectively. The first term describes the diffusion (how the colony spreads in space), while the second term is the logistic growth rate describing the proliferation of the bacteria.

We also wanted to describe the concentrations of mRNAs and proteins inside the cell in the growing colony. $a_i$ stands for the concentration of the $i$th species inside the cell. In a growing colony – without any reactions between the chemical species – $ca$ is a conserved quantity, so it should fulfil the continuity equation ($\partial_0 ca_i = -\boldsymbol{\nabla} \cdot \vec{j}_{ca_i}$). Furthermore, the reaction of the intracellular components can be taken into account as well:

$$\partial_0 ca_i = -\boldsymbol{\nabla} \cdot \vec{j}_{ca_i} + cf_i(\vec{a}), \tag{11}$$

where $\vec{j}_{ca_i}$ is the flux of $ca_i$ and $f_i$ describes the reactions of $a_i$. For the reactions the same model (Table S2) was used as before. The flux of $a_i$ is proportional with the flux of the cells ($\vec{j}_c$) – since the content of the cells can move only with the cells – and can be calculated as:

$$\vec{j}_{ca_i} = a_i \vec{j}_c. \tag{12}$$

##### 2.3.1 Dimensionless form

To reduce the number of the parameters, a dimensionless form was used. The unit of the cell concentration was expressed with the carrying capacity and the distance was given based on the diffusion coefficient:

$$M_c = [F], \quad m = \left[\sqrt{\frac{D}{k_4}}\right], \tag{13}$$

which results $\hat{F} = 1$, $\hat{\mathrm{D}} = 1$, and the dimensionless model:

$$\partial_0 \hat{c} = \nabla^2 \hat{c} + \hat{k}\hat{c}(1 - \hat{c}), \quad \partial_0(\hat{c} \cdot \hat{a}_i) = \boldsymbol{\nabla} \cdot \hat{a}_i \boldsymbol{\nabla} \hat{c} + \hat{c} f_i(\vec{\hat{a}}), \tag{14}$$

with the following parameters and concentrations:

$$\hat{L} = \frac{L}{\sqrt{\frac{D}{k_4}}}, \quad \hat{T}_s = T_s k_4, \quad \hat{k} = \frac{kF}{k_4}, \quad \hat{c} = \frac{c}{F} \tag{15}$$

where $L$ is the size of the simulation domain and $T_s$ is the length of the simulation.

##### 2.3.2 Simulation details

The PDEs were solved on a $200 \times 200$ grid with the finite–difference method. The time integration was fulfilled with the Euler method [10] for diffusion and with the fully composite Patankar–Euler method [7] for the stochastic reaction kinetics equations with a $\hat{dt} = 10^{-3}$ time step. The domain size and the length of the simulation were $\hat{L}_x = \hat{L}_y = 500$, $\hat{T}_s = 10$. For the Laplace approximation a nine point stencil was used [11] to be able to observe a completely circular growing colony. For the boundary, a periodic boundary condition was used. The simulations were written in C++ using Eigen and OpenMP for parallelization, while the visualization was fulfilled in MATLAB.

In the experiments the cells were kept in a dark environment before the external forcing started. Based on this experimental condition, the initial condition in the simulations (for the concentration of the mRNA ($m_i$) and proteins ($P_i$)) was set according to the fixed point of the ODE system (Table S2) with zero light intensity ($\hat{I} = 0$). These initial conditions were applied in every grid point, and their value can be found in Table S5. For the initial concentration of the cells, a short simulation was performed first to get a smooth Gaussian–like concentration distribution. Briefly, the cells were placed in the middle of the simulation domain within a radius $\hat{R}_0 = 5$ and with an initial concentration $\hat{c}_0 = 1$. After that, a short simulation ($\hat{T}_0 = 10^{-1}$) was performed – with the same parameters and method described before –, where only the cell diffusion was taken into account ($\partial_0 \hat{c} = \nabla^2 \hat{c}$). The output of this short simulation was used as the initial condition for the cells in the colony growth simulations.

The cells do not remain active forever. In the center of the colony, cells enter stationary phase, their metabolism slows down and the oscillations eventually stop. To reflect this behaviour and to confine the reactions to a reasonable cell concentration, two threshold concentrations ($\hat{c}_{min} = 10^{-6}$, $\hat{c}_{max} = 0.99$) were introduced. A given reaction rate ($\hat{r}$) can be modified with these threshold values: $\hat{r}\prime = \hat{r}\Theta(\hat{c}_{min} - \hat{c})\Theta(\hat{c} - \hat{c}_{max})$, where $\Theta$ is the Heaviside step function. For the logistic growth of the cells only the lower, while for the reactions of the mRNAs and proteins both the lower and the upper thresholds were applied.

##### 2.3.3 The effect of noise

The simulations are sensitive to the strength of the noise. To understand this effect, we have carried out simulations with different system sizes ($\hat{\Omega}$). In Figure S4 the first line corresponds to the deterministic case (infinite system size, or zero noise), while the noise strength is increasing with the rows. In column $a$ – with constant light intensity – the pattern gets less and less symmetric, the concentric rings become less circular. However, in the case of the resonance (column $b$), due to the synchronization the rings remain circular, and the effect of the noise is smaller.

There is another interesting phenomenon in the case of period-2 ($c$), period-4 ($d$) and chaos ($e$). In the deterministic simulations (row $A$) one can clearly see the high and low intensity peaks in period doubling ($Ac$) or the peaks with different intensities and radii in the case of chaos ($Ae$). While the intensity difference between the peaks are decreasing in period doubling, the rings remain circular. However, in the case of the chaotic pattern, the rings become less and less circular with the noise, because a chaotic system is sensitive even to small changes and with a positive Lyapunov exponent the distance between the solutions increases exponentially. In the stochastic simulations the external light source synchronises the cells, except in the chaotic patterns, where a strong desynchronization can be observed. Interestingly, in every non-autonomous case ($b - e$), with strong noise (row $E$) we can observe regular, circular patterns and the rings have the same intensity. It is a known phenomena that noise can stabilize a chaotic system [12, 13]. Here we can observe the stabilizing effect of noise as the chaotic ring pattern ($Ef$) becomes a regular pattern ($Ee$).

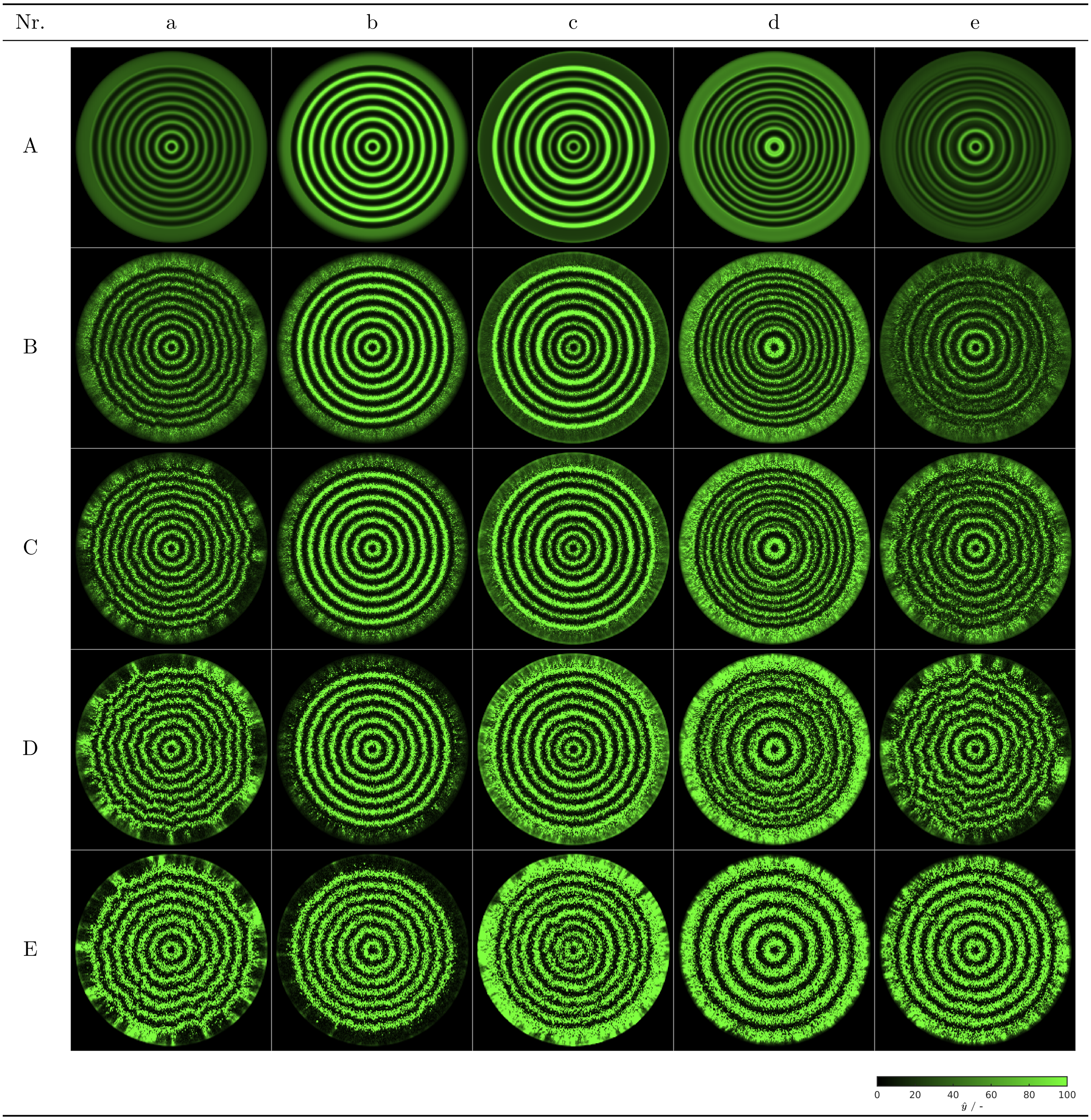


**Figure S4:** The effect of the noise. The columns of the figure corresponds to Figure 4 of the main text and to Figure S5: *a*: constant light with ($\hat{I} = 30\%$), *b*: resonance ($\hat{T} = 13$), *c*: period two ($\hat{T} = 25$), *d*: period four ($\hat{T} = 18$), *d*: chaos ($\hat{T} = 7.45$). While the noise is increasing with the rows of the figure (as the volume is decreasing) *A*: $\hat{\Omega} = \infty$ (deterministic), *B*: $\hat{\Omega} = 5$, *C*: $\hat{\Omega} = 2$, *D*: $\hat{\Omega} = 1$, *E*: $\hat{\Omega} = 0.5$. Corresponding movies of the simulations are provided as supplementary movies.

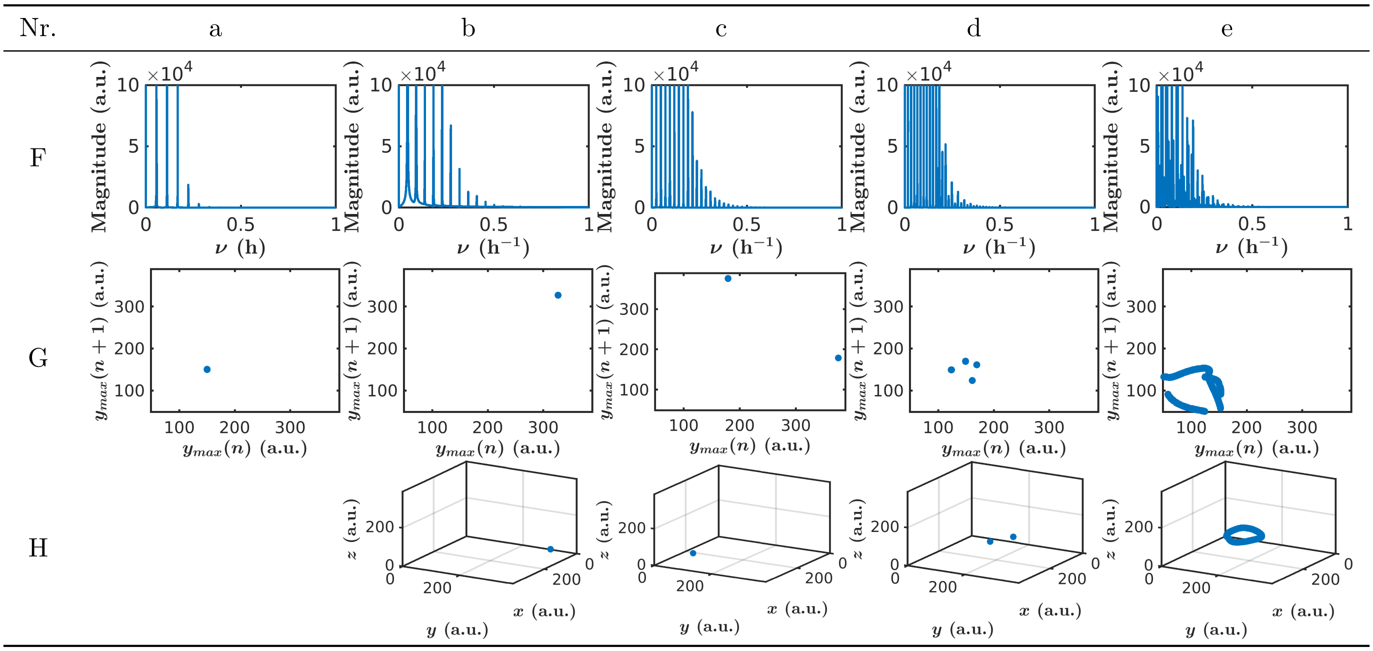


**Figure S5:** Additional analyses complementing Figure 4 from the main text. *F*: the Fourier spectra, *G*: maxima return map, H: stroboscopic map. (In the case of the continuous light – column *a* – the system is autonomous so we do not show the stroboscopic map.) *a*: constant light with ($\hat{I} = 30\%$), *b*: resonance ($\hat{T} = 13$), *c*: period two ($\hat{T} = 25$), *d*: period four ($\hat{T} = 18$), *d*: chaos ($\hat{T} = 7.45$)

**Table S5:** The initial condition in the simulations. The concentrations were calculated from the fixed point of the non-dimensional ODE system (Table S2) with zero light intensity ($\hat{I} = 0$, dark state). ($\hat{r}_{2,i} = [\hat{m}_i] - \left[\hat{P}_i\right]$ results $[\hat{m}_i]^\star = \left[\hat{P}_i\right]^\star$ in the fixed point.)

| Parameter | Value | Parameter | Value | Parameter | Value |
|---|---|---|---|---|---|
| $[\hat{m}_x]_0$ | 6.38 | $[\hat{m}_y]_0$ | 116.82 | $[\hat{m}_z]_0$ | 5.01 |
| $[\hat{P}_x]_0$ | 6.38 | $[\hat{P}_y]_0$ | 116.82 | $[\hat{P}_z]_0$ | 5.01 |

### Processing the colony images

#### Calculating the fluorescent intensity as a function of the radius

At the end of the experiments, we took microscopic images of the colonies. We then converted the images into radius–fluorescence functions, because this way it is easier to see the desynchronization speed and to compare different colonies. After finding the center of the colony (the ,,mass center” of the highest fluorescent intensity region), the distance of every pixel from the center (their radius) was determined. In the figures we show both the original data and the smoothed version (moving average with $10^4$ points) for the individual colonies and for their average.

#### Space–time transformation

To be able to calculate the time period of the oscillator and to compare the phase of the outer forcing (blue light source) and the oscillator, we transformed the space–fluorescence curves into time–fluorescence curves. In Figure S6a we followed the size of a growing colony. The colony growth was described with the following equation:

$$r = g(t) = k_2 - k_3 \exp(-t/k_1), \tag{16}$$

where $k_1$, $k_2$ and $k_3$ were fitted parameters and their value can be found in Table S6. The details of the fitting procedure can be found in section 2.2.

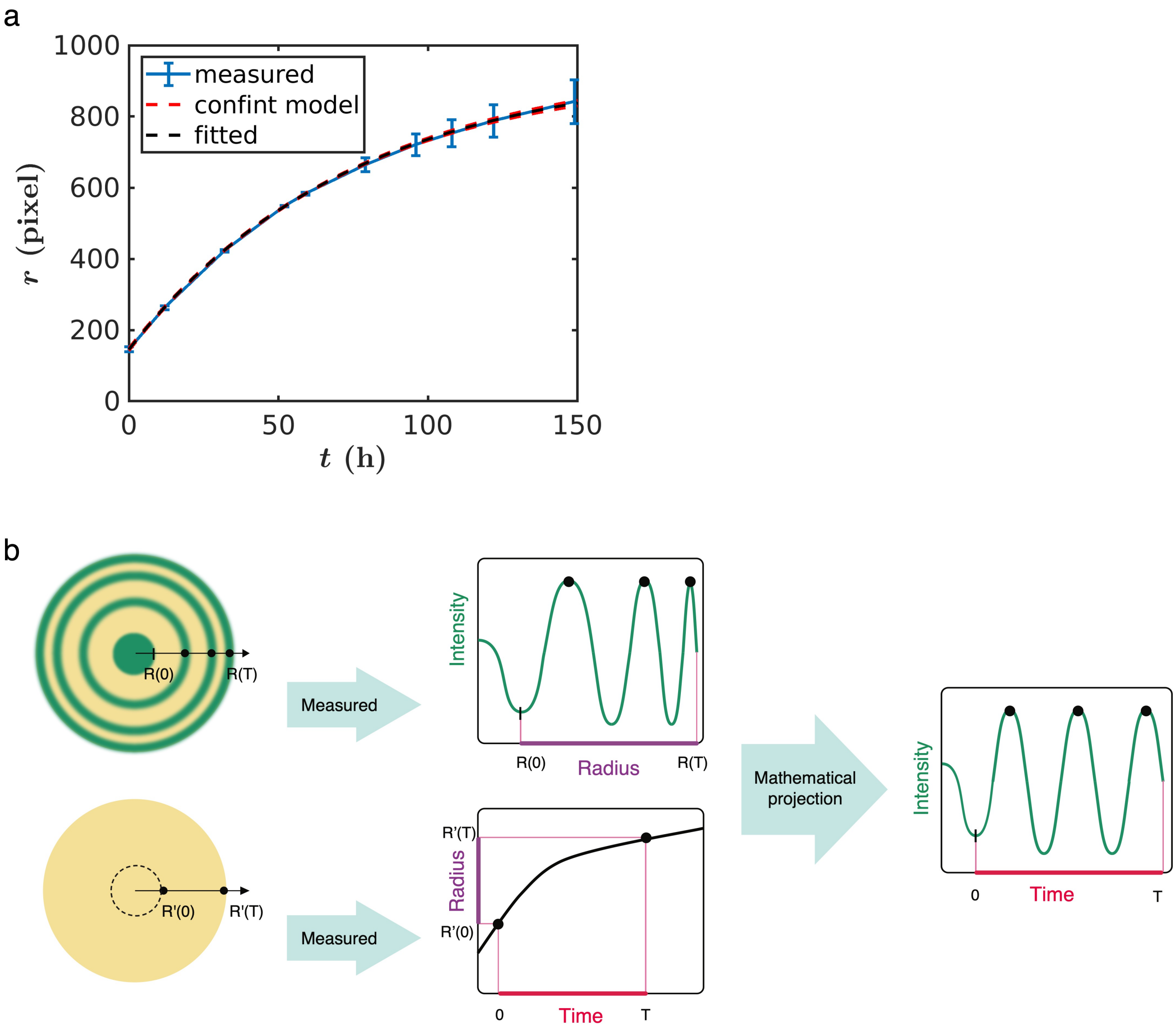


**Figure S6:** Conversion of radius to time. a) Change of the radius ($r$) of colonies as a function of growth time ($t$) with 100 % constant light intensity. The measured data show the mean of three biological replicates (blue line), with error bars depicting the s.d. The fitted model (dashed black line) has 95 % confidence interval for the fitted function (red dashed line). b) Schematic representation of the main steps for image processing of the colonies.

In the space–time transformation, the initial time is calculated based on the initial size of the colony after starting from a single cell and 20 h of incubation at 21 $°C$. To make sure that at the beginning of the experiment we get time zero after the transformation (in spite of the small experimental differences in the initial colony size), we included an additional linear transformation ($h$). With this transformation, the initial and the final sizes of the colony will give zero hours and the length of the experiment, respectively ($h(R_T) = g(T)$, $h(R_0) = g(0)$). The linear transformation is the following:

$$r' = h(r) = \frac{r - R_0}{R_T - R_0}(g(T) - g(0)) + g(0), \tag{17}$$

where $r'$ is the new radius, $r$ is the original radius, $R_0$ is the initial colony size, $R_T$ is the colony size at the end of the experiment and $t$ is the length of the experiment. We carried out this transformation in the case of the modulated external forcing (where the phase of the signal was important), while for the experiments with constant light intensities, we carried out this adjustment only with the final size of the colony.

The space–time transformation was carried out the following way:

$$t' = g^{-1}(h(r)), \tag{18}$$

first the linear transformation ($h$) for the space (to make sure, that the initial and final time will be 0 and $T$), and after that the space–time transformation, which is the inverse function of the fitted curve ($g^{-1}$). The sketch of this transformation can be seen in Figure S6b.

**Table S6:** The fitted parameters (and the 95 % confidence interval of the fitting) for the colony growth experiments. The value of $k_1$ is given in hours, while $k_2$ and $k_3$ are given in pixels.

| Parameter | Value | Parameter | Value | Parameter | Value |
|---|---|---|---|---|---|
| $k_1$ | $74.4 \pm 2.7$ | $k_2$ | $944 \pm 17$ | $k_3$ | $799 \pm 16$ |
